# In-silico studies of Neurocognitive and Neuropharmacological effect of Bacopa monnieri (L.)

**DOI:** 10.1101/2021.01.20.427542

**Authors:** Satyam Sangeet, Arshad Khan

**Affiliations:** Indian Institute of Science Education and Research (IISER), Bhopal, Madhya Pradesh, India; National Institute of Technology (NIT), Arunachal Pradesh, India

**Keywords:** Neurotrophins, *Bacopa monnieri*, Docking, Molecular simulations, ADME, Phytochemicals

## Abstract

Different Indian therapeutic plants have picked up consideration for their restorative properties against neurodevelopmental disorders lately. *Bacopa monnieri (B. monnieri)*, being one of them, finds its utilization significantly in the treatment of cognition and learning. Despite the fact that it possesses such great capacity to treat neurological issues, how precisely it confers its influence is muddled. This study makes a stride towards knowing which phytochemical is significantly associated with granting *B. monnieri* with therapeutic properties. The docking investigation and the molecular simulation studies proposes that there is no single phytochemical included that imparts *B. monnieri* a significant medicinal effect. It is really the blend of dominant part of natural phytochemicals present in *B. monnieri* that bestows the anti-neurological activity to *B. monnieri.* The ADME studies shows the convergence of properties of phytochemicals of *B. monnieri* with that of commercially available drugs which suggests that phytochemicals of *B. monnieri* can used as a potential drug source to treat neurodegenerative and neurodevelopmental disorders.

## INTRODUCTION

Neurodegenerative disorders (NDs) are recognized to be noteworthy threats to human health. Numerous symptoms are associated with NDs including memory loss, tremors, forgetfulness, agitation. Many kinds of mechanism lead to NDs such as apoptosis, protein aggregation, oxidative stress, excitotoxicity and ageing. NDs constitute various types of disorders which include Parkinson’s^1^, Huntington’s^2^ Alzheimer’s disease and other dementias^3^, amyotrophic lateral sclerosis^4^, frontotemporal dementia^5^, spinocerebellar ataxias^6^, stroke^7^, meningitis^8^, encephalitis^9^, tetanus^10^, epilepsy^11^, multiple sclerosis^12^, motor neuron disease^13^, migraine^14^, tension-type headache^15^, medication overuse headache^16^, brain and nervous system cancers^17^, and a residual category of other neurological disorders. There is huge burden of NDs on the global, regional and national levels. Vast number of people have been affected by these diseases. Global burden of NDs are estimated to be around 450 million^18^.

The NDs have a major involvement of Neurotrophins, which are group of proteins having related structures and functions and have an important involvement in survival ^19, 20^, maintenance ^21^, function^22, 23^ regulation^24^ and development^25^ of neurons. The mammalian cells include four types of Neurotrophins: Brain-derived Neurotrophic Factor (BDNF)^26^, Neurotrophin 4 (NT 4/5)^27^, Nerve Growth Factor (NGF)^28^ and Neurotrophin 3 (NT-3)^29^. Neurotrophins belong to a class of growth factors hence named as neurotrophic factors^30^. Neurotrophic factors that mammalian cells secrete plays an important role in the survival of neurons by preventing them from participating in programmed cell death. Decrease in the level of Neurotrophins results in several neurodegenerative disorder^31, 32^. Several studies have considered Neurotrophins to be a potential target for the treatment of neurological disorders ^33, 34, 35^.

All the four Neurotrophins bind individually, by same affinity with their common receptor P75^NTR 36^ and a specific trk receptor kinase, and thus activates the signal cascade response. p75^NTR^ receptor is structurally related to the tumour necrosis factor receptors (p55^TNFR^ and p75^TNFR^). In NGF, residues Lys32, Lys34 and Lys95 have been found to be actively associated with the p75NTR binding. Also, the involvement of residues Trp21, Asp30, Ile31, Glu35, Lys88, Arg100 and Arg103 with the binding of p75NTR has been reported^37^. Several synthetic drugs are being used to treat different neurological disorders^38^, for example, Multiple Sclerosis (MS) a neurological disease is treated by wide arrays of repurposed drugs. Some drugs which proved to be effective in mitigating the effect of MS are mitoxantrone^39^, cyclophosphamide^40^, cladribine^41^, amiloride^42^ and ibudilast^43^. Use of synthetic drugs is being restricted because of their various kind of side effects such as headache, pain, toxicity, nausea, alopecia, male and women infertility and risk of malignancy. Mitoxantrone is reported to impart many kinds of side effects in the patient suffering from MS^44^. Cyclophosphamide and cladribine also have shown significant side effects^45^.

Various components of medicinal plants are known to have therapeutic effects pertaining to neurodegenerative disorders^46^.It is revealed that different metabolites present in plants such as phenolics, isoprenoids and alkaloids etc. are mainly responsible for neurotherapeutic actions. Several traditional plants such as Indian traditional plants (Plants of Ayurveda), Chinese traditional plants, Utopian traditional plants and Persian traditional plants etc. have been reported widely to heal brain disorders and diseases of nervous system. Some example of the traditional plants used against neurodegenerative disorders are *Curcuma longa* L.^47^, *Murraya koenigii* (L.) Spreng^48^, *Cassia obtisufolia* L.^49^, *Sanguisorba officinalis* L.^50^, *Rosmarinus officinalis* L. (Rosemary)^51^, *Huperzia serrata* (Thunb.) Trevis. (Qiancengta)^52^, *Paeonia suffruticosa*^53^, *Melissa officinalis* L.^54^.

Phytochemicals from medicinal plants have been exploited for their potential in targeting neurodegenerative disorders by regulating the Neurotrophins. Example of some plants whose phytochemicals target NDs by affecting Neurotrophins are *Aster scaber*^55^, *Camellia sinensis*^56^, *Curcuma longa*^57^, *Ginkgo biloba* (L)^58^, *Liriope platyphylla*^59^ and *Magnolia officinalis*^60^.

*Bacopa monnieri* (BM) is a significant Ayurvedic medicinal plants that has been in use since ancient time to enhance brain function and to improve intelligence as well as memory^61^. Role of *B. monnieri* in cognitive and memory enhancement has been studied extensively^62^. *B. monnieri* contains many bioactive components such as saponins, triterpenoid bacosaponins and saponin glycosides^62^. Stough et al^63^ reported that *B. monnieri* extract enhanced the cognition safely and effectively. Standard extracts of *B. monnieri* have shown improvement in many neurodegenerative disorders owing to presence of different bioactive phytochemicals^60,63,64,65,66^

Treatment of neurodegenerative disorders by targeting Neurotrophins using the plant extract of *Bacopa monnieri* (BM) has not been reported yet. Lal & Lal^67^ has reported phytochemicals present in the plant extract of BM using GC-MS study. Even, with the advancement in technology, it is still not clear as to how BM imparts its effect on the ND. Also, the molecular pathway behind the action is not clearly understood.

Thus, the current study focuses on the individual activity of *B. monnieri* phytochemicals against the Neurotrophins by studying its molecular docking analysis and molecular simulations. This study will provide us with an insight as to which phytochemical is majorly involved in imparting *B. monnieri* its medicinal property and whether the neuroprotective activity of *B. monnieri* is due to a single phytochemical or a cumulative effect of all the phytochemicals combined.

## MATERIALS AND METHOD

### Ligand and Receptor Preparation

The approved drugs for the treatment of Multiple Sclerosis (Cladribine, Cyclophosphamide and Mitoxantrone) were chosen as the standard drugs^38^. Based on the literature evidences^67^, the structure of 26 phytochemicals of *B. monnieri* (Table 1) were downloaded from PubChem (https://pubchem.ncbi.nlm.nih.gov/) database^68^. The phytochemicals were converted from 2D to 3D by using Open Babel software^69^ and were energy minimized using Chem3D^70^ software using MM2 minimize.

**Table 1.**
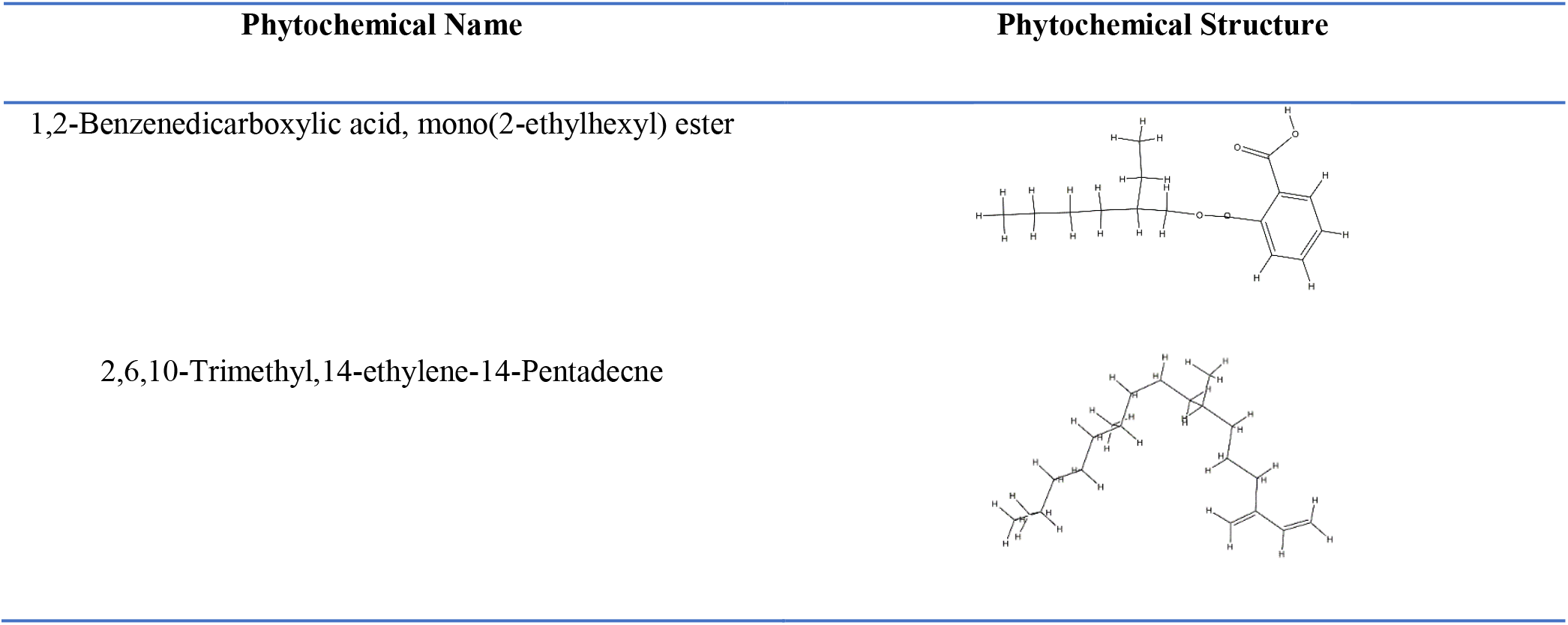
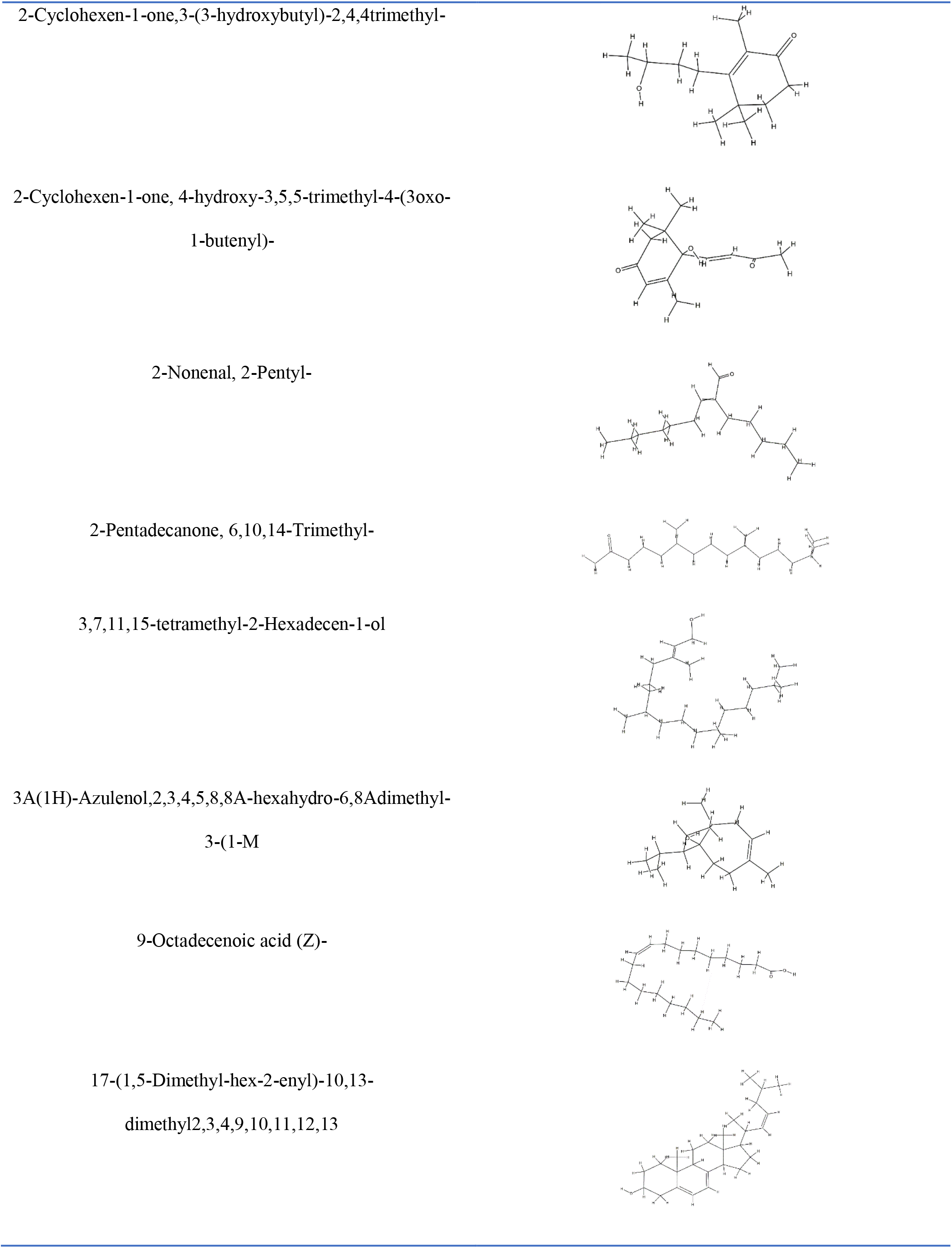
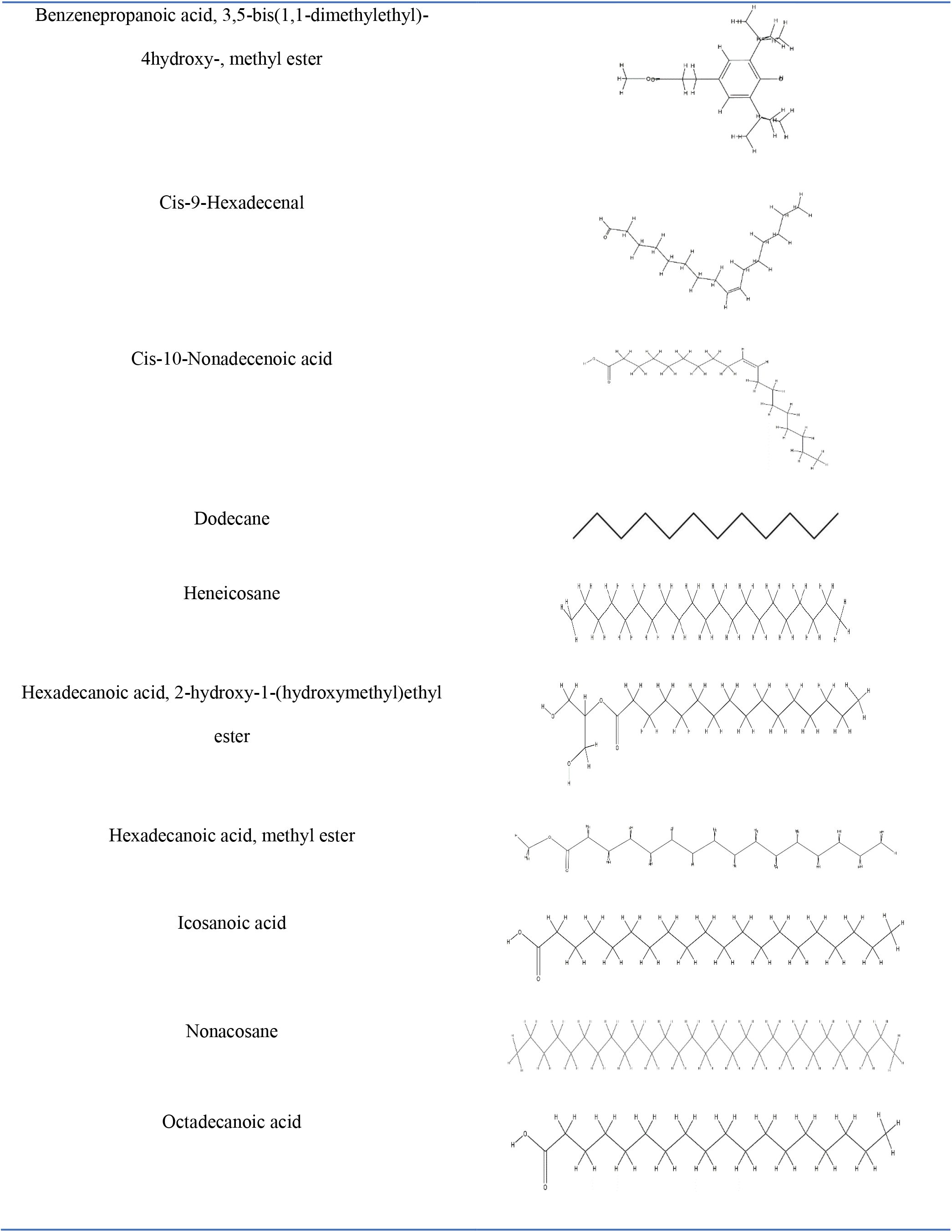
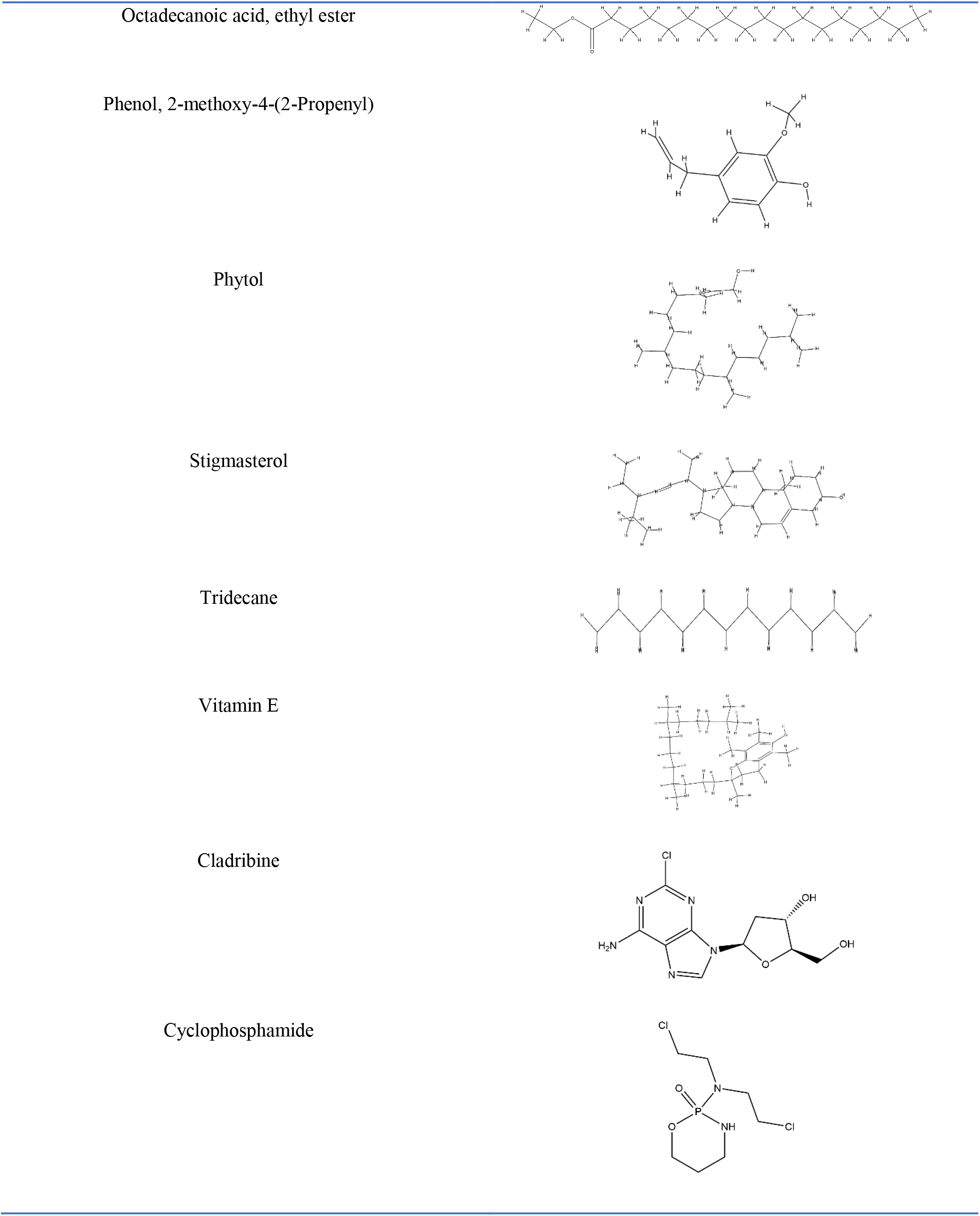
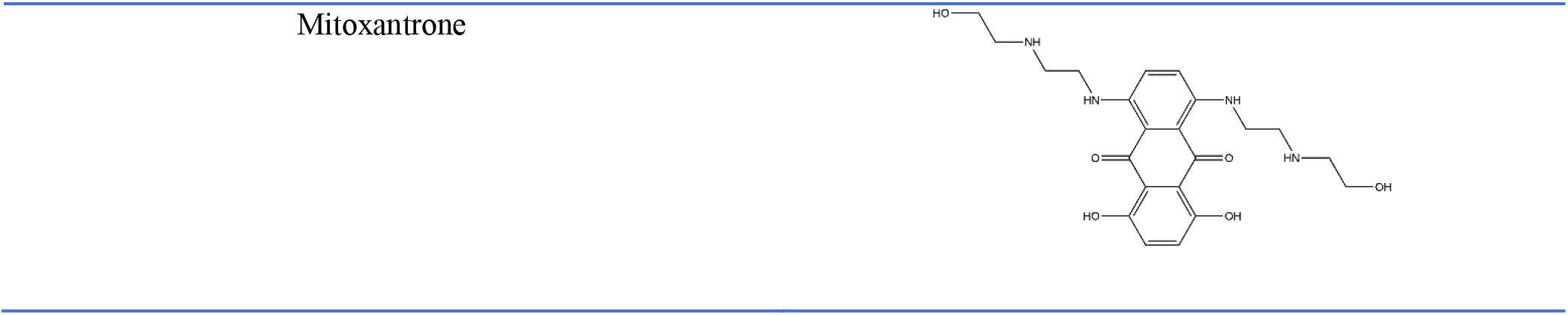
Phytochemicals and commercial drugs and their structure

Receptor PDB structures of Brain-Derived Neurotrophic Factor (PDB ID: 1BND), Neurotrophin 3 (PDB ID: 1B8K), Neurotrophin 4 (PDB ID: 1B98) and Nerve Growth Factor (PDB ID: 4EDX) were downloaded from Protein Data Bank (www.rcsb.org). The receptors were prepared by deleting all the water molecules and unwanted residues and energy minimizing it using Chimera software^71^.

### Molecular Docking

PatchDock online server (https://bioinfo3d.cs.tau.ac.il/PatchDock/) was employed for the molecular docking studies between the Neurotrophins and the phytochemicals. The clustering RMSD values were set at default (4.0 Å) as suggested by the server. The respective Neurotrophin structures were uploaded in the Receptor molecule option and the phytochemical structures were uploaded in the Ligand Molecule option. The PatchDock score and the ACE value were tabulated and the result was visualized in Discovery studio^72^.

### Molecular Simulation

The online server CABS-flex 2.0 (http://biocomp.chem.uw.edu.pl/CABSflex2) was utilized for the molecular simulations of the docked results. The values were set as the default parameter as suggested by the server. The results obtained were downloaded in the form of excel sheet. The Root Mean Square Fluctuation (RMSF) curves were visualized using Jupyter Notebook and matplotlib python package.

### ADME Studies

Adsorption, Digestion, Metabolism, Excretion and Toxicity studies were performed using SWISSADME server (http://www.swissadme.ch; ^73^). The structure of drugs and phytochemicals were uploaded and the corresponding drug likeness features such as lipophilicity, solubility and drug likeness score were obtained.

## RESULTS

The molecular docking between the Neurotrophins and the phytochemicals of *Bacopa monnieri* (*B. monnieri*) was carried out. Literature studies^68^ were looked for creating the phytochemical database. Once the docking process was finished the molecular simulations were carried out to find the RMSF values of the involved amino acids followed by the ADME studies to see the drug likeness of the phytochemicals.

### Computational Docking

Out of the 26 phytochemicals docked against the four Neurotrophins, only two of them showed interactions with BDNF, two of them showed interaction with Neurotrophin 3, 17 of them showed interaction with Neurotrophin 4 and 11 of them showed interactions with NGF (Table 2).

**Table 2.**
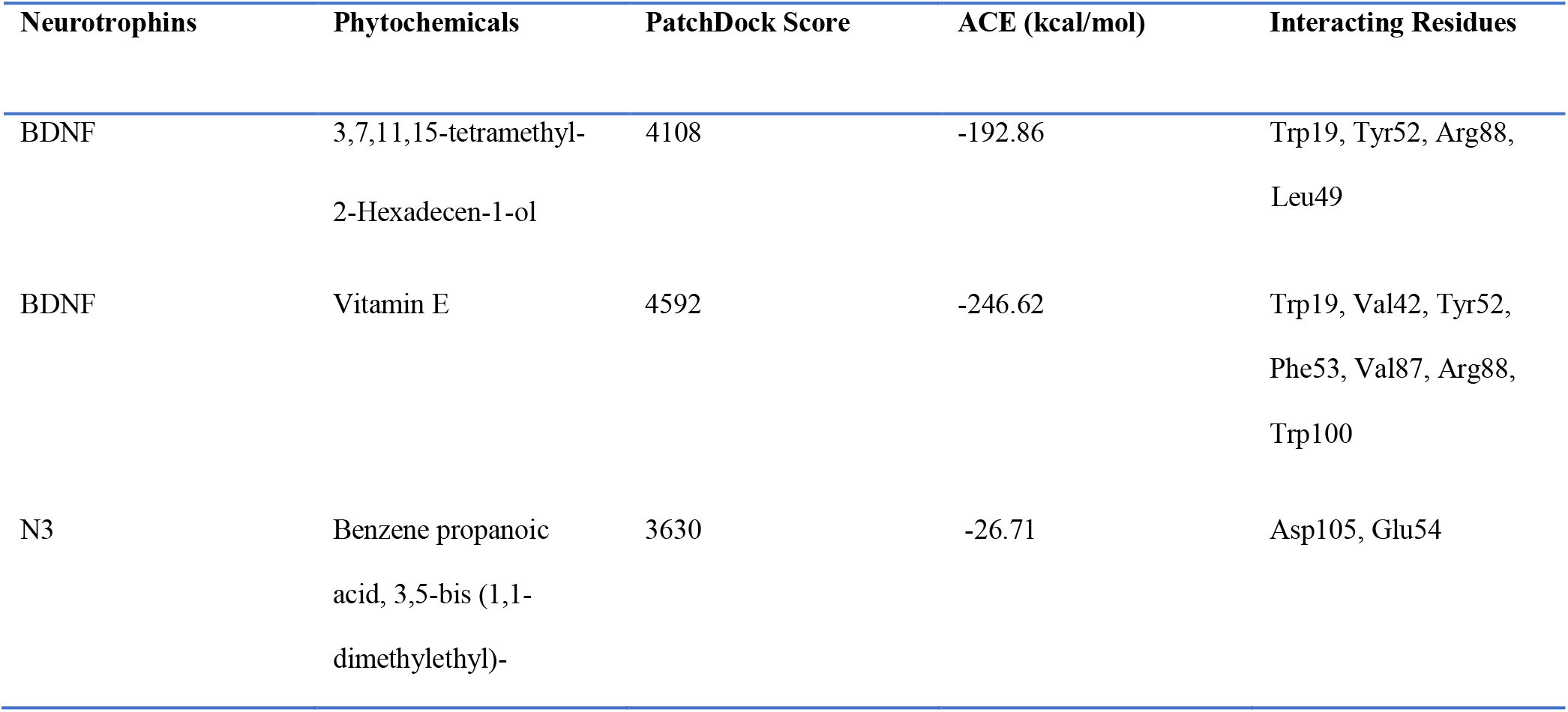
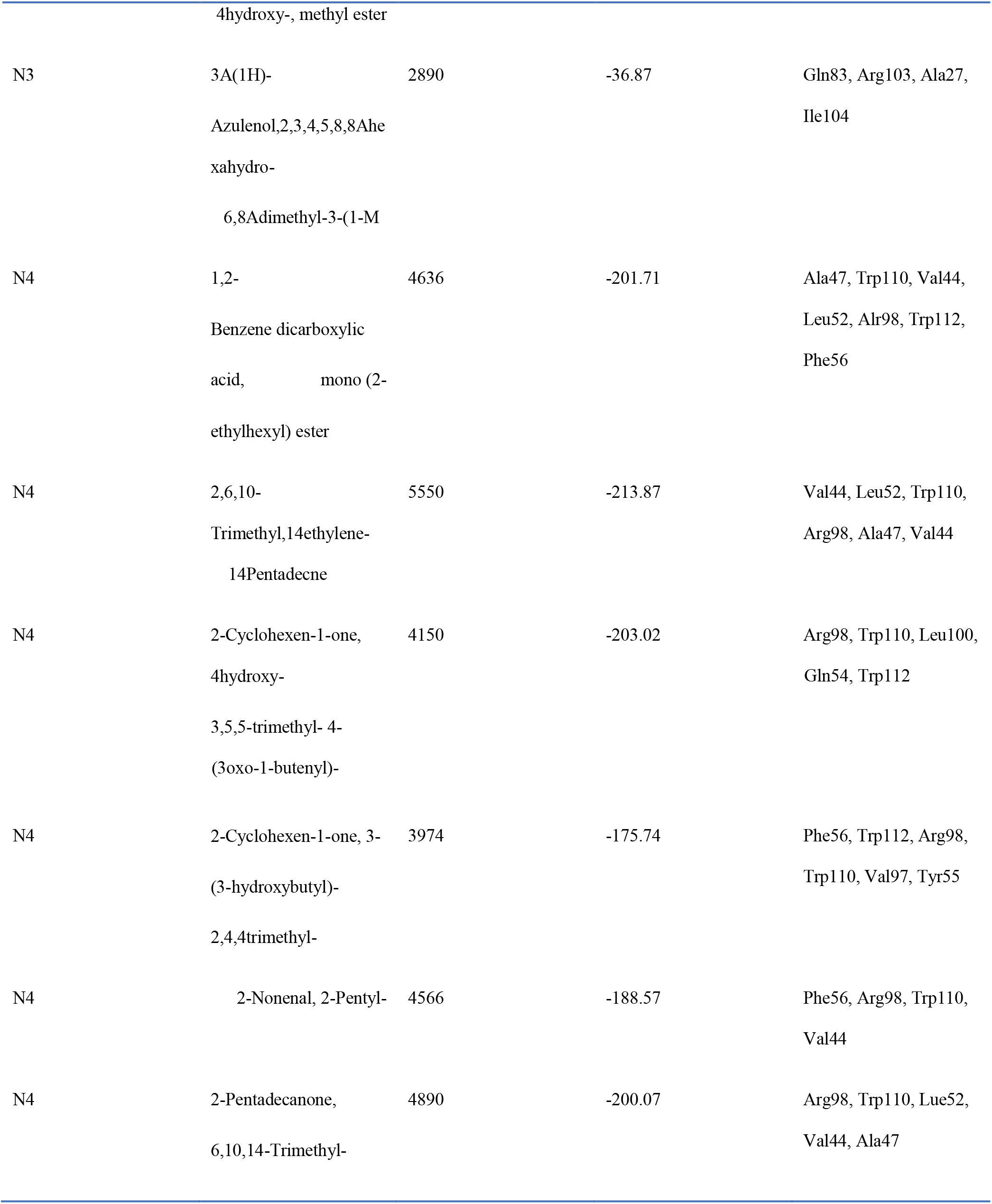
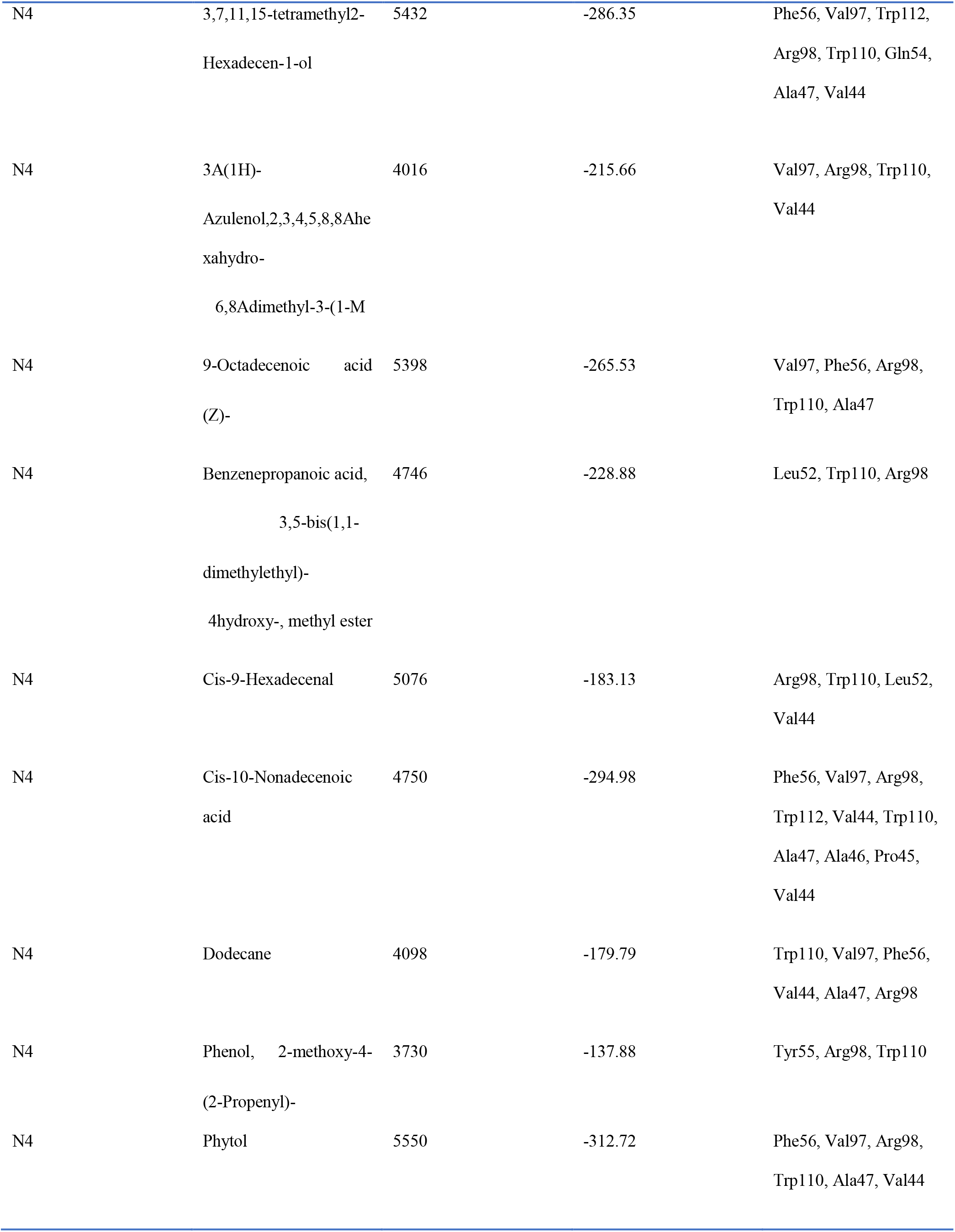
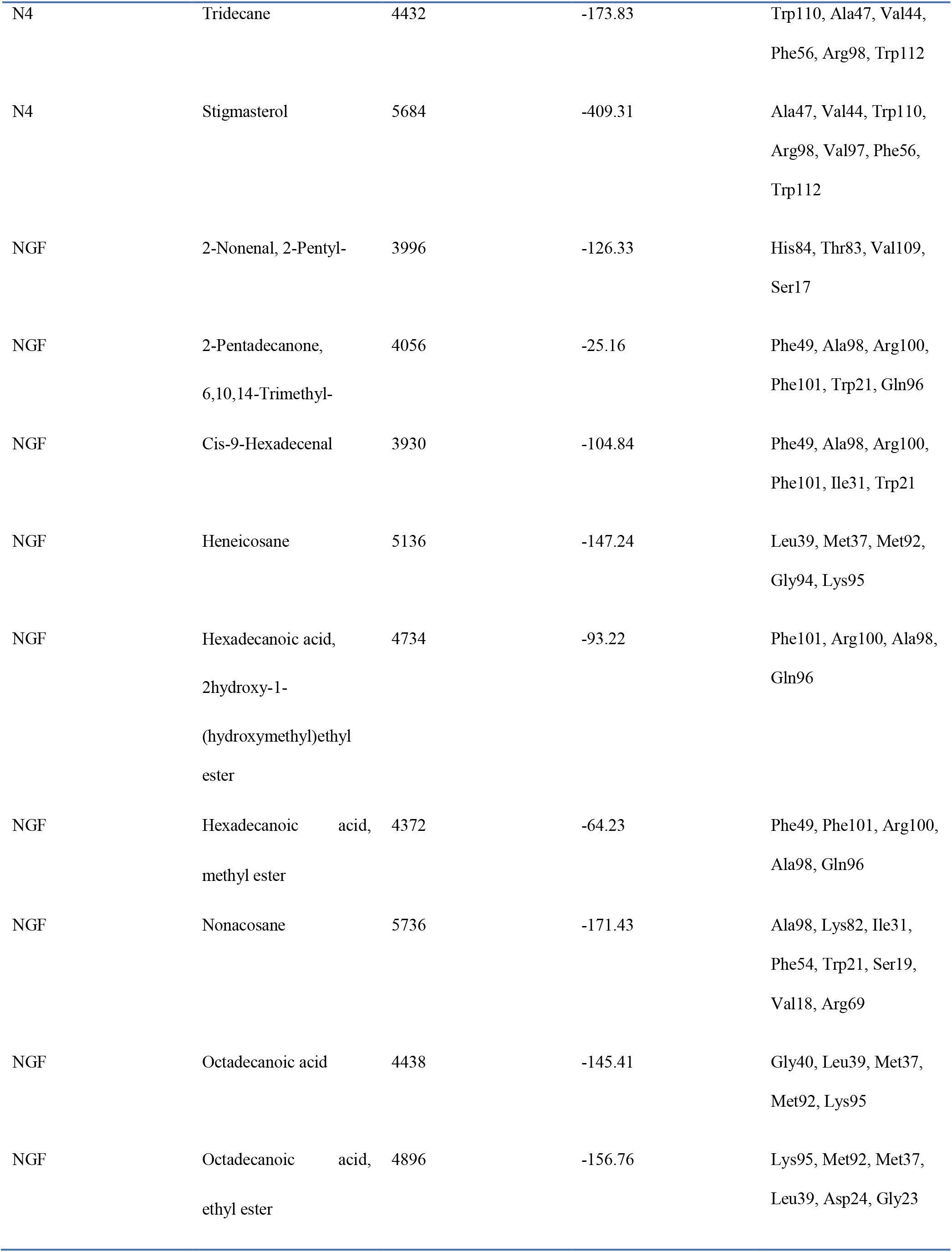
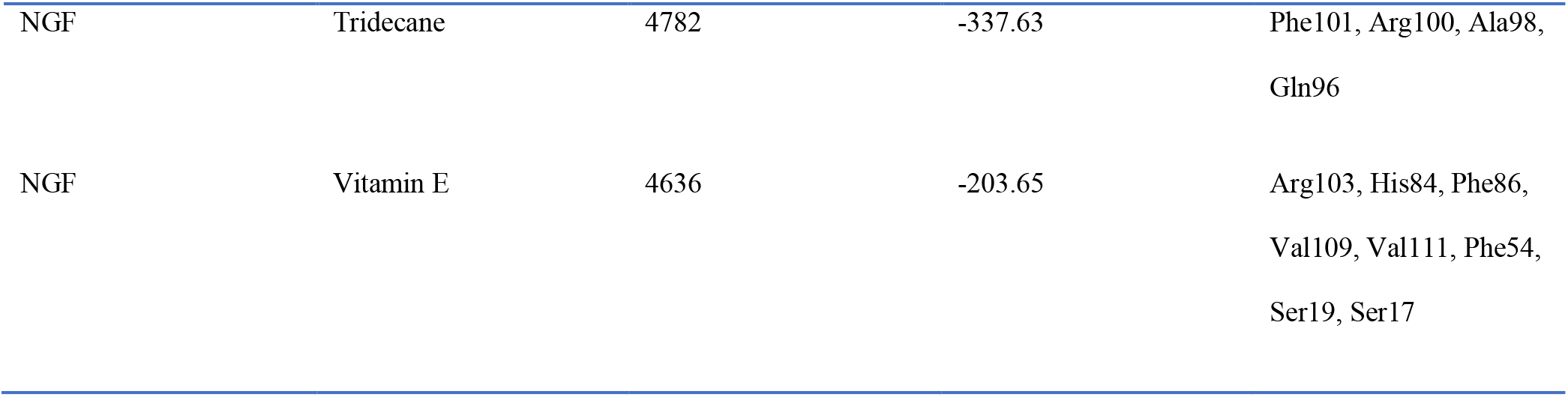
Neurotrophins along with the interacting phytochemicals and the respective PatchDock Score, ACE value and the interacting amino acid residues.

### Brain – Derived Neurotrophic factor (BDNF)

BDNF naturally interacts with p75^NTR^ with residues Gln84, Arg104, Asp30, Arg101, Arg88, Asp93, Lys95, Arg97 and Lys96^36^. Docking between BDNF and the phytochemicals of *Bacopa monnieri* showed that only two of the 26 natural phytochemicals show a feasible interaction (Table 2). 3,7,11,15-tetramethyl2Hexadecen-1-ol showed hydrogen bonding with Arg88 and hydrophobic interactions with Trp19, Leu49 and Tyr52 with a docking score of 4,108 and an ACE value of −192.86 kcal/mol (Figure 1a). Vitamin E showed hydrophobic interactions with Trp19, Val42, Tyr52, Phe53, Val87, Arg88 and Trp100 with a docking score of 4,592 and an ACE of −246.62 kcal/mol (Figure 1b). Thus, this investigation shows that when the phytochemicals of *Bacopa monnieri* is used as an anti-inflammatory agent against the neurodegenerative disorders, the interaction of phytochemicals with BDNF shows the involvement of two phytochemicals: 3,7,11,15-tetramethyl-2-Hexadecen-1-ol and Vitamin E.

**Figure 1.**
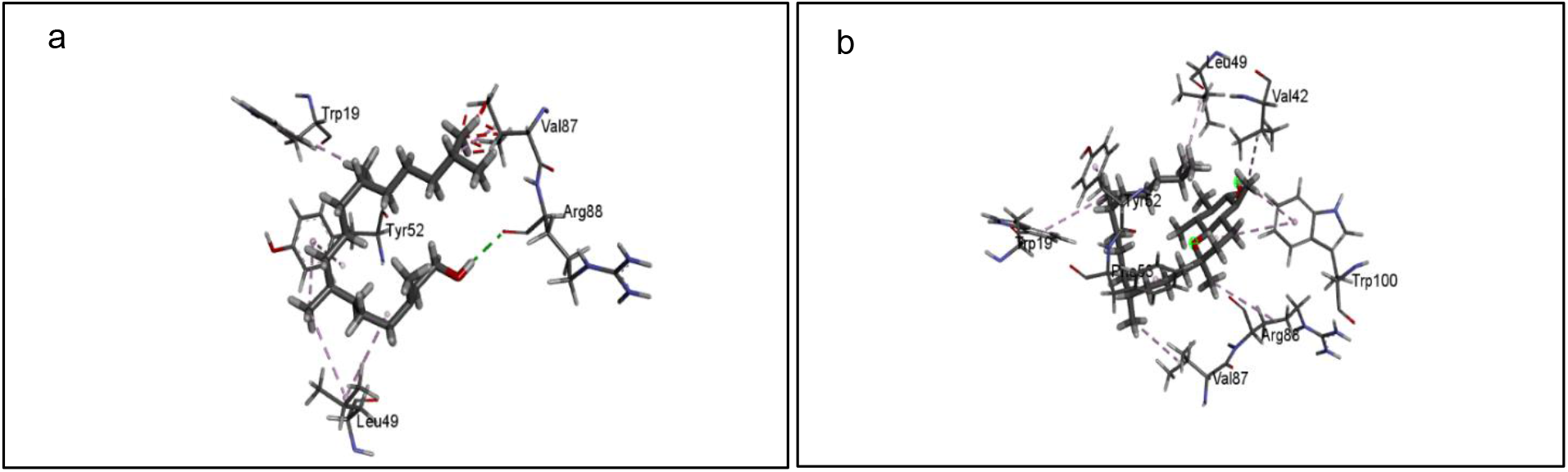
Docking interactions between BDNF and (a) 3,7,11,15-tetramethyl-2-Hexadecen-1-ol (b) Vitamin E

### Neurotrophin 3 (N3)

Neurotrophin 3 interacts with p75NTR through residues Arg103, Gln83, Asp29, Arg87, His33, Arg31, Arg100, Glu92, Leu96 and Lys95^36^. Two phytochemicals from *B. monnieri* interacted with Neurotrophin 3 which are shown in Table 2. Benzene propanoic acid, 3,5-bis(1,1-dimethylethyl)-4hydroxy-, methyl ester showed hydrogen bonding with Asp105 and a carbon hydrogen bonding with Glu54 with a docking score of 3,630 and an ACE value of −26.71 kcal/mol (Figure 2a). 3A(1H)-Azulenol,2,3,4,5,8,8Ahexahydro6,8Adimethyl-3-(1-M showed hydrogen bonding with Gln83 an Arg103. It also showed hydrophobic interactions with Ala27 and Ile104 with a docking score of 2,890 and an ACE value of −36.87 kcal/mol (Figure 2b).

**Figure 2.**
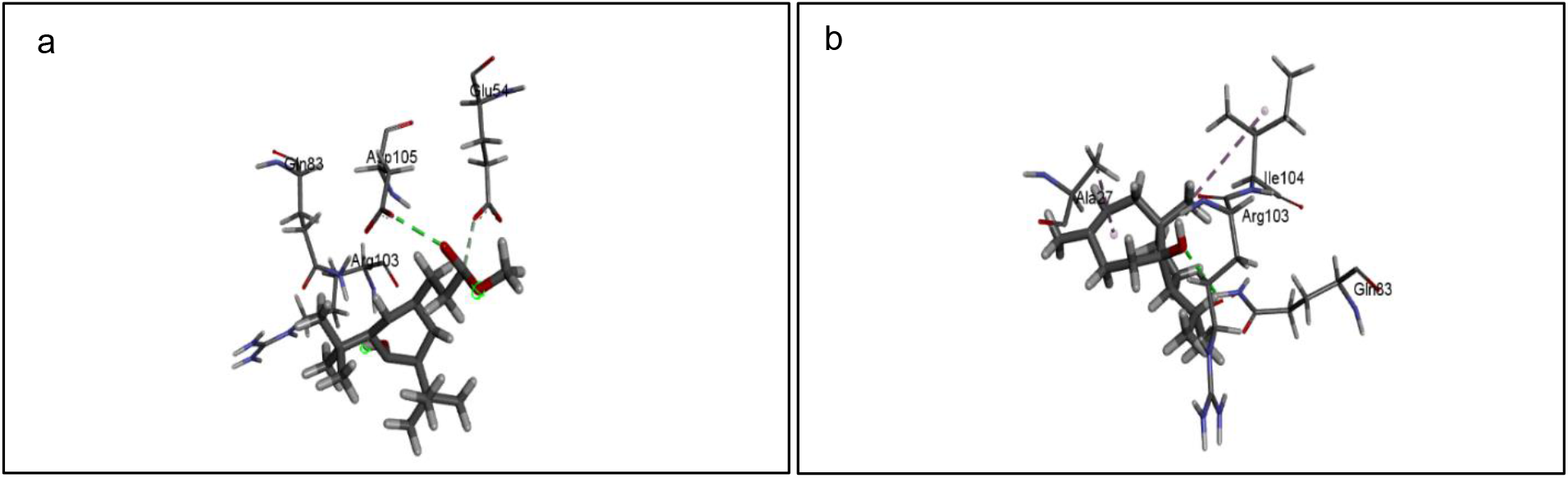
Docking interactions between Neurotrophin 3 (N3) and (a) Benzene propanoic acid, 3,5bis(1,1dimethylethyl)-4hydroxy-, methyl ester (b) 3A(1H)-Azulenol,2,3,4,5,8,8A-hexahydro-6,8Adimethyl-3(1-M

### Neurotrophin 4 (N4)

Neurotrophin 4 interacts with p75NTR through residues Gln83, Arg114, Tyr85, Asp32, Arg98, Arg34, Arg111, Asp75 and Arg107^36^. 17 phytochemicals interacted with the Neurotrophin 4. The interaction of the phytochemicals can be found in Table 3. 2,6,10-Trimethyl,14-ethylene-14-Pentadecne showed hydrophobic interactions with Val44, Ala47, Arg98, Leu52 and Trp110 with a docking score of 5,550 and an ACE of −213.87 kcal/mol (Figure 3a). 3,7,11,15-tetramethyl-2-Hexadecen-1-ol showed carbon hydrogen bonding with Gln54 and hydrophobic interactions with Arg98, Ala47, Val97, Val44, Trp110, Trp112 and Phe56 with a docking score of 5,432 and an ACE value of −286.35 kcal/mol (Figure 3b). 9-Octadecenoic acid (Z)- showed hydrogen bonding with Trp110 and hydrophobic interactions with Arg98, Ala47, Val97, Trp110 and Phe56 with a docking score of 5,398 and an ACE of −265.53 kcal/mol (Figure 3c). Phytol showed hydrophobic interactions with Val44, Arg98, Ala47, Val97, Phe56 and Trp110 with a docking score of 5,550 and an ACE of −312.72 kcal/mol (Figure 3d). Stigmasterol showed hydrophobic interactions with Trp110, Val44, Ala47, Arg98, Val97, Phe56, Trp110 and Trp112 with a docking score of 5,684 and an ACE of 409.31 kcal/mol (Figure 3e).

**Table 3.**
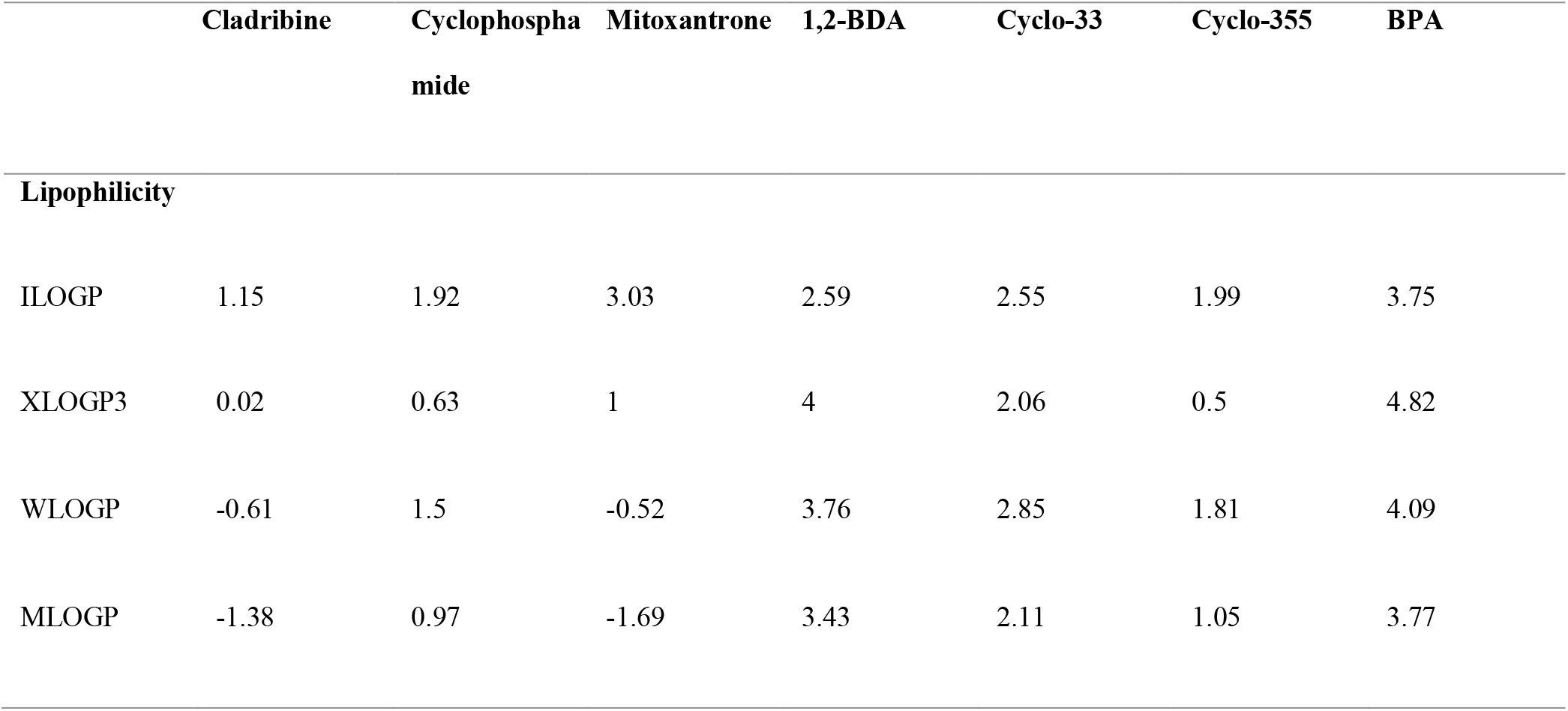
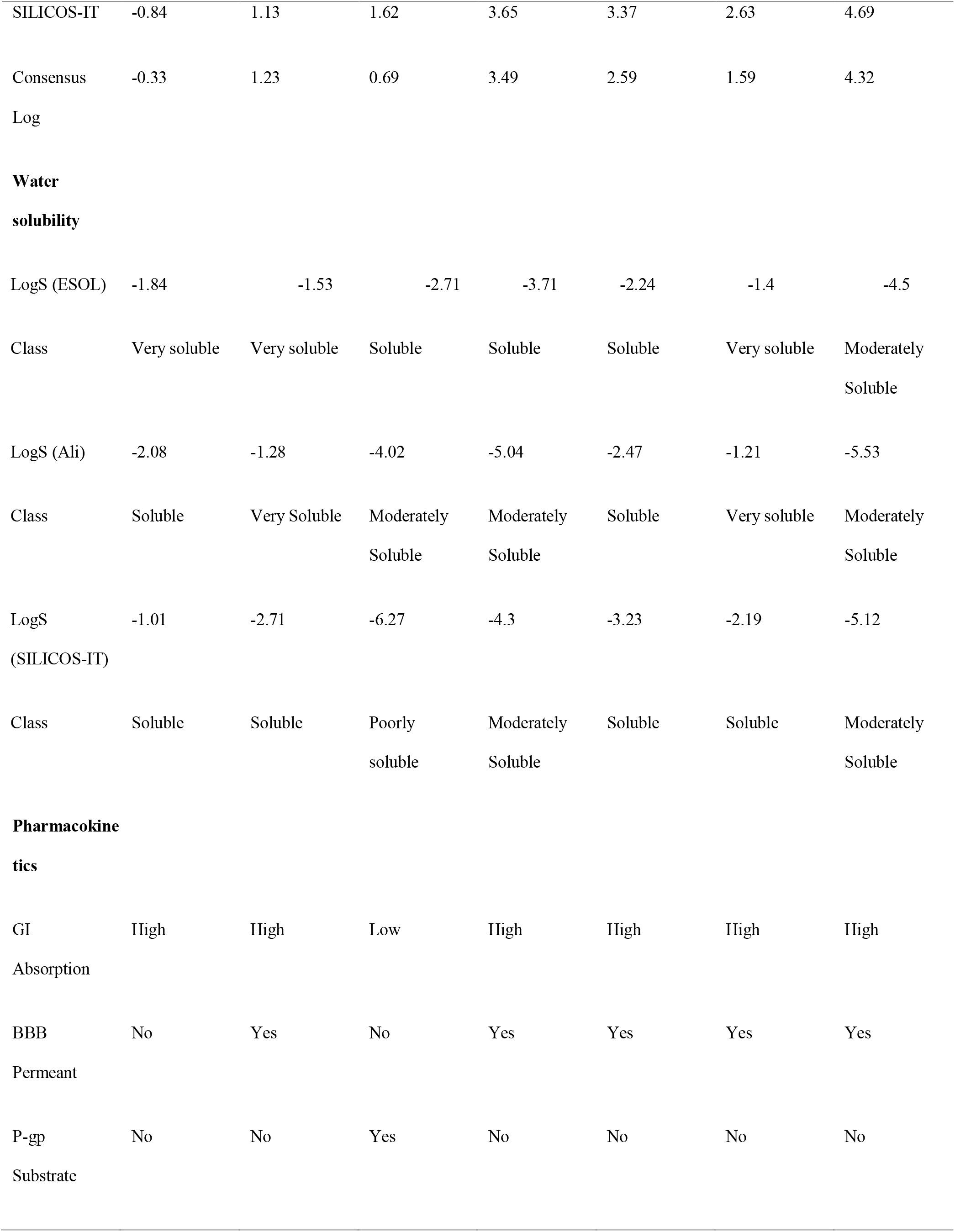
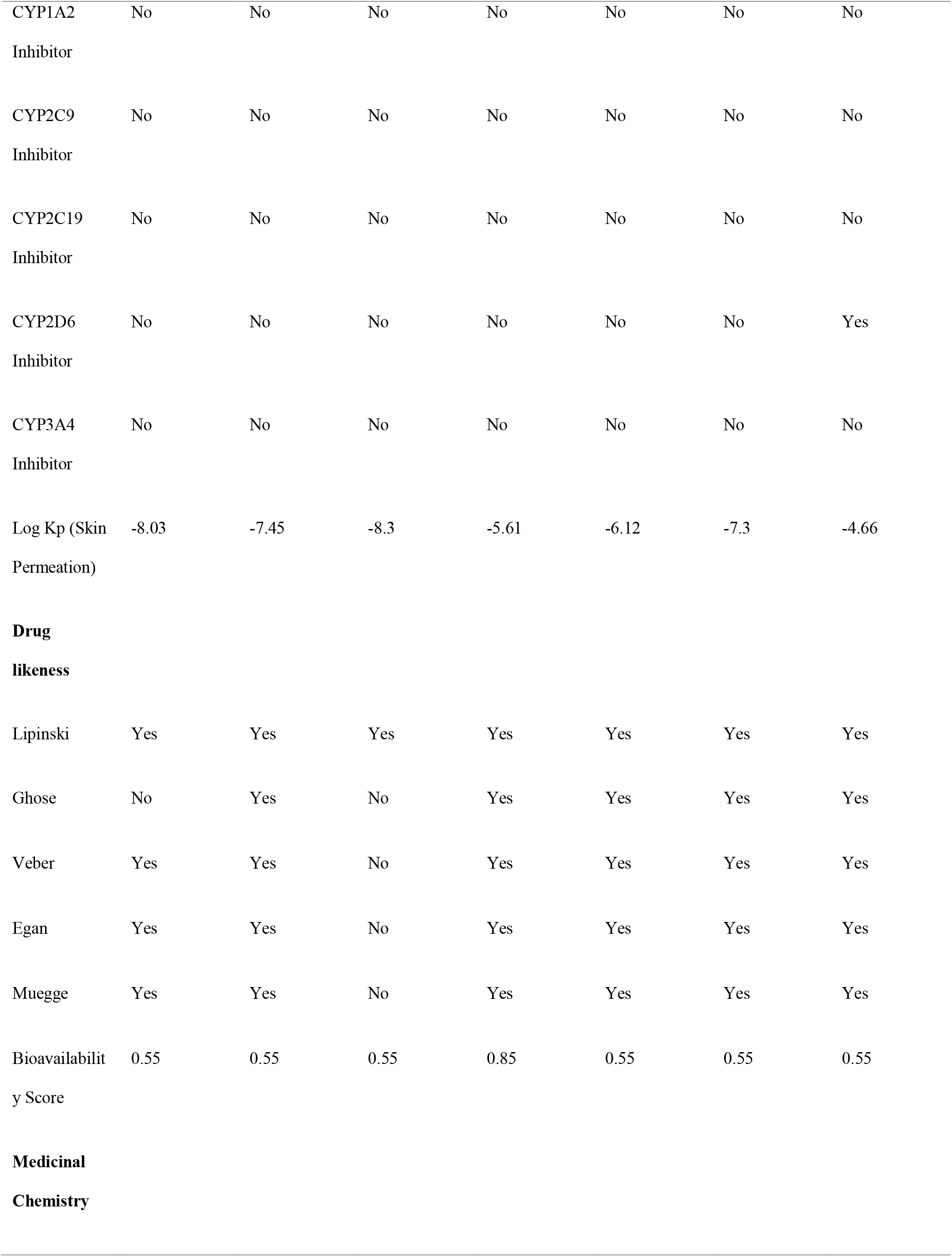
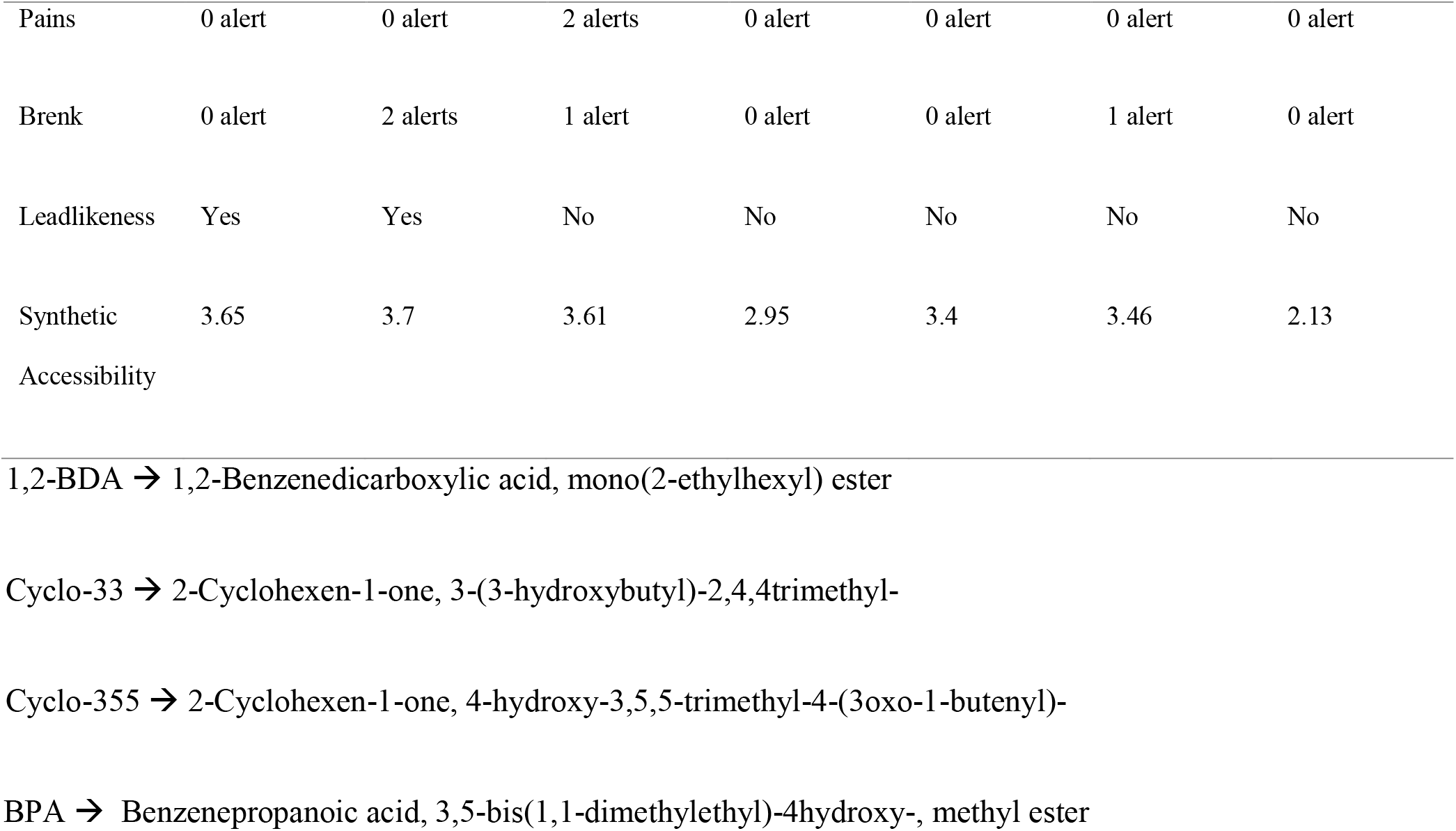
ADME results for the commercial drugs and screened phytochemicals

**Figure 3.**
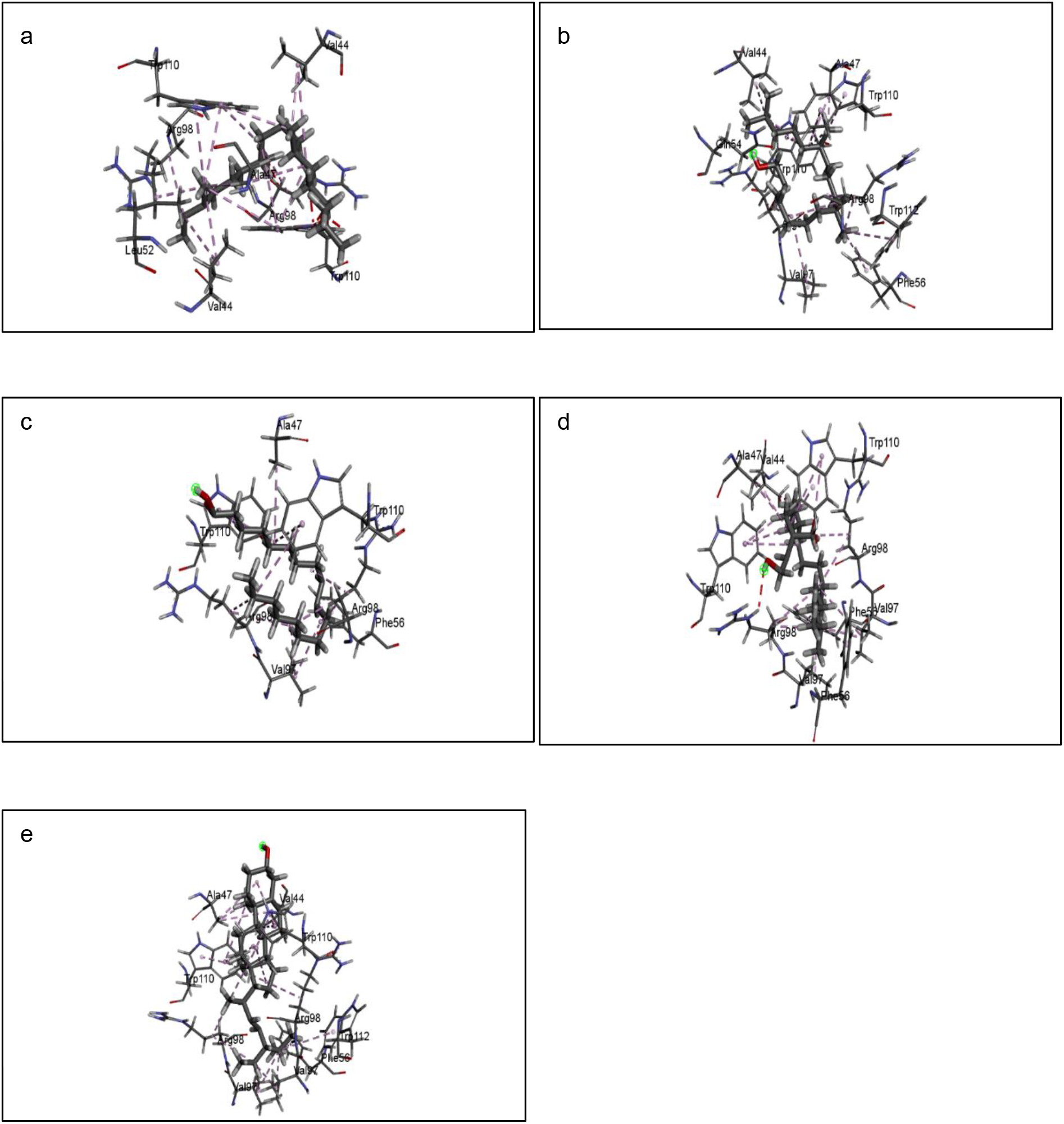
Docking interactions between Neurotrophin 4 and (a) 2,6,10-Trimethyl,14-ethylene-14Pentadecne (b) 3,7,11,15-tetramethyl-2-Hexadecen-1-ol (c) 9-Octadecenoic acid (Z)- (d) Phytol (e) Stigmasterol

### Nerve Growth Factor (NGF)

NGF interacts with p75NTR through residues Lys32, Lys34, Lys95, Arg103, His84, Arg100, Glu35, Lys88, Asp30, Asp93, Trp21 and Ile31^36^. 11 phytochemicals interacted with Nerve Growth Factor (NGF). The interaction between phytochemicals and the NGF is shown in Table 4. Heneicosane and Nonacosane showed better results as compared with other phytochemicals. Heneicosane showed hydrophobic interactions with Met37, Leu39, Met92 and Lys95 with a docking score of 5,136 and an ACE of −147.24 kcal/mol (Figure 4a). Nonacosane showed hydrophobic interactions with Lys32, Ala98, Ile31, Arg59, Trp21 and Phe54 with a docking score of 5,736 and an ACE value of −171.43 kcal/mol (Figure 4b).

**Figure 4.**
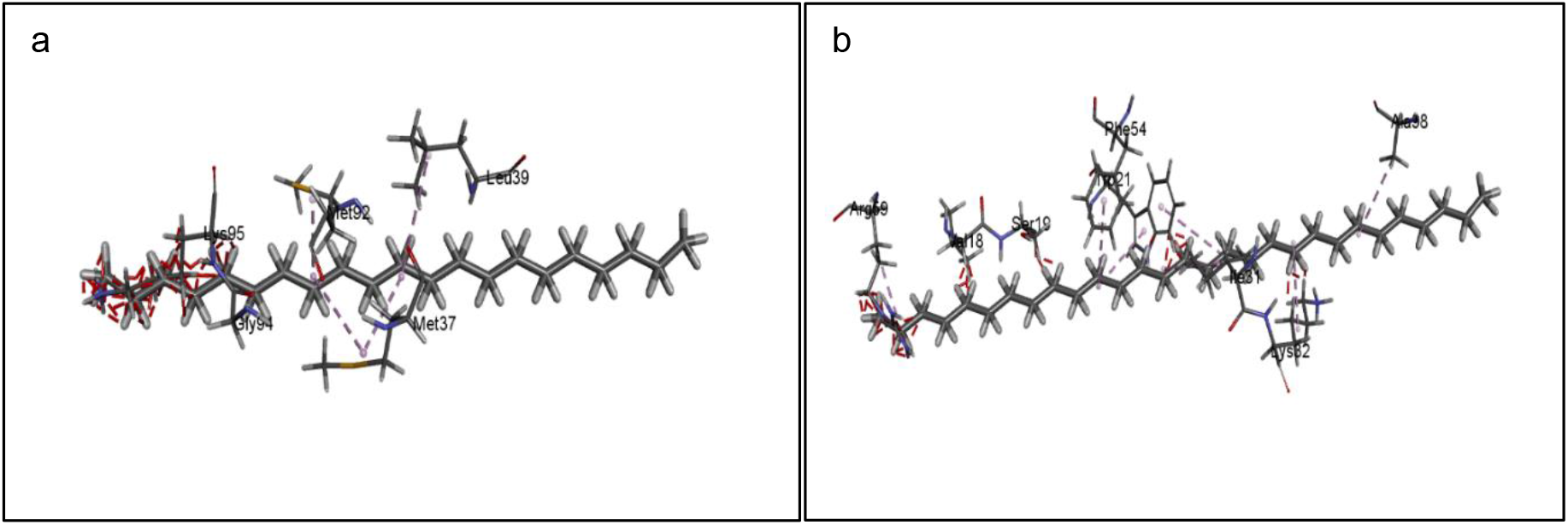
Docking interactions between Nerve Growth Factor (NGF) and (a) Heneicosane (b) Nonacosane

### Molecular Simulations

Molecular simulations of the docking studies were performed for each Neurotrophin using CABS-Flex 2.0 online software (http://biocomp.chem.uw.edu.pl/CABSflex2/index). The corresponding RMSF values obtained were then plotted to see the individual fluctuations of the amino acid actively participating and interacting with the respective Neurotrophins.

### Brain Derived Neurotrophic Factor (BDNF)

The molecular simulation studies of Vitamin E (Figure 5c) suggest that the curve almost fits closely to that of BDNF along with its natural ligand. Thus, from the curve it can be extrapolated that Vitamin E can possess the potential of imparting the anti-neurodegenerative activity to *B. monnieri*.

**Figure 5.**
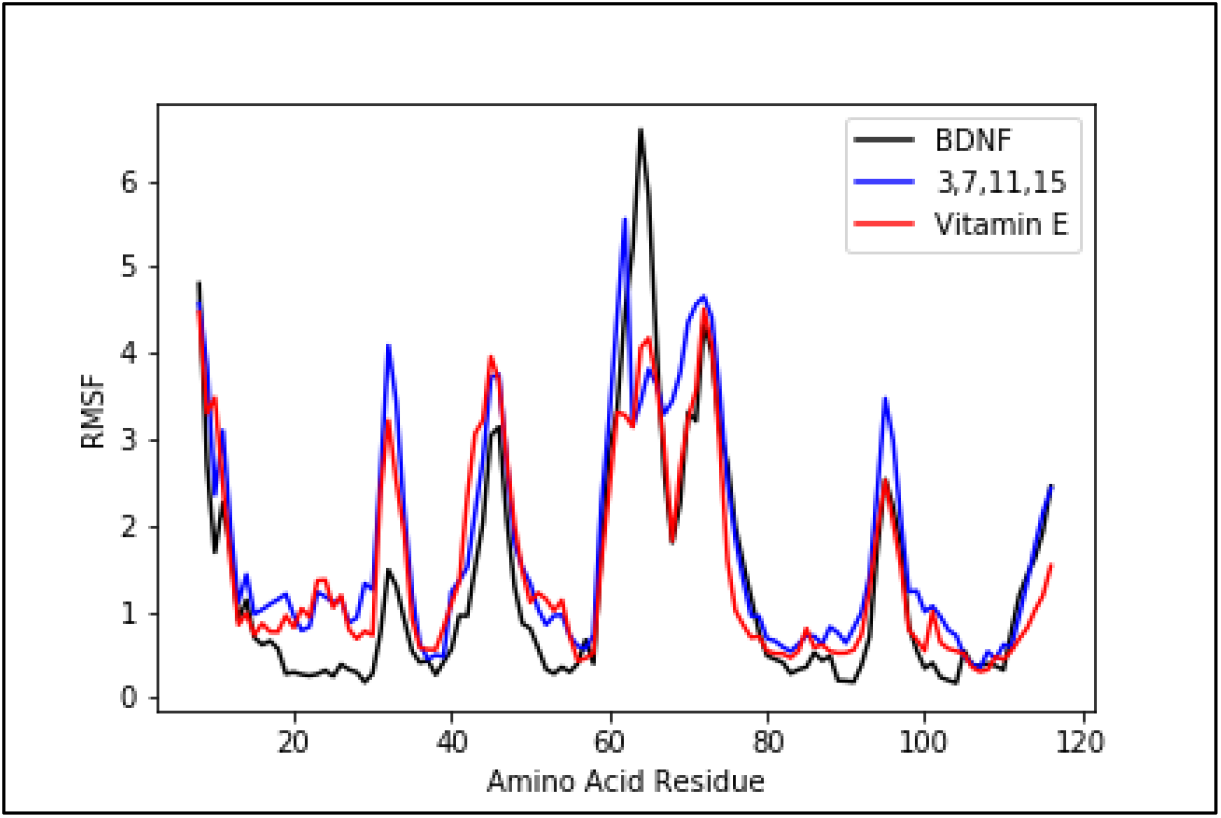
Molecular Simulation studies of BDNF docked with natural ligand, 3,7,11,15tetramethyl-2-Hexadecen-1-ol and Vitamin E

### Neurotrophin 3 (N3)

Looking at the molecular simulation curve of N3 with the respective phytochemicals it can be seen that Benzene propanoic acid, 3,5-bis(1,1-dimethylethyl)-4hydroxy-, methyl ester (Figure 6b BPA) shows better results as compared with 3A(1H)-Azulenol,2,3,4,5,8,8A-hexahydro-6,8Adimethyl-3-(1-M (Figure 6c 3A(1H)). When the active amino acid is looked upon on the X-axis, it can be seen that the BPA curve for Gln83 has a dip which goes beyond the curve of N3 suggesting that BPA is more stable than N3. A similar result can be seen for Arg103 as well. Comparing the molecular simulation results with the PatchDock score obtained it can be confirmed that BPA might also provide anti-depressant and anti-neurodegenerative activity to *B. monnieri*.

**Figure 6.**
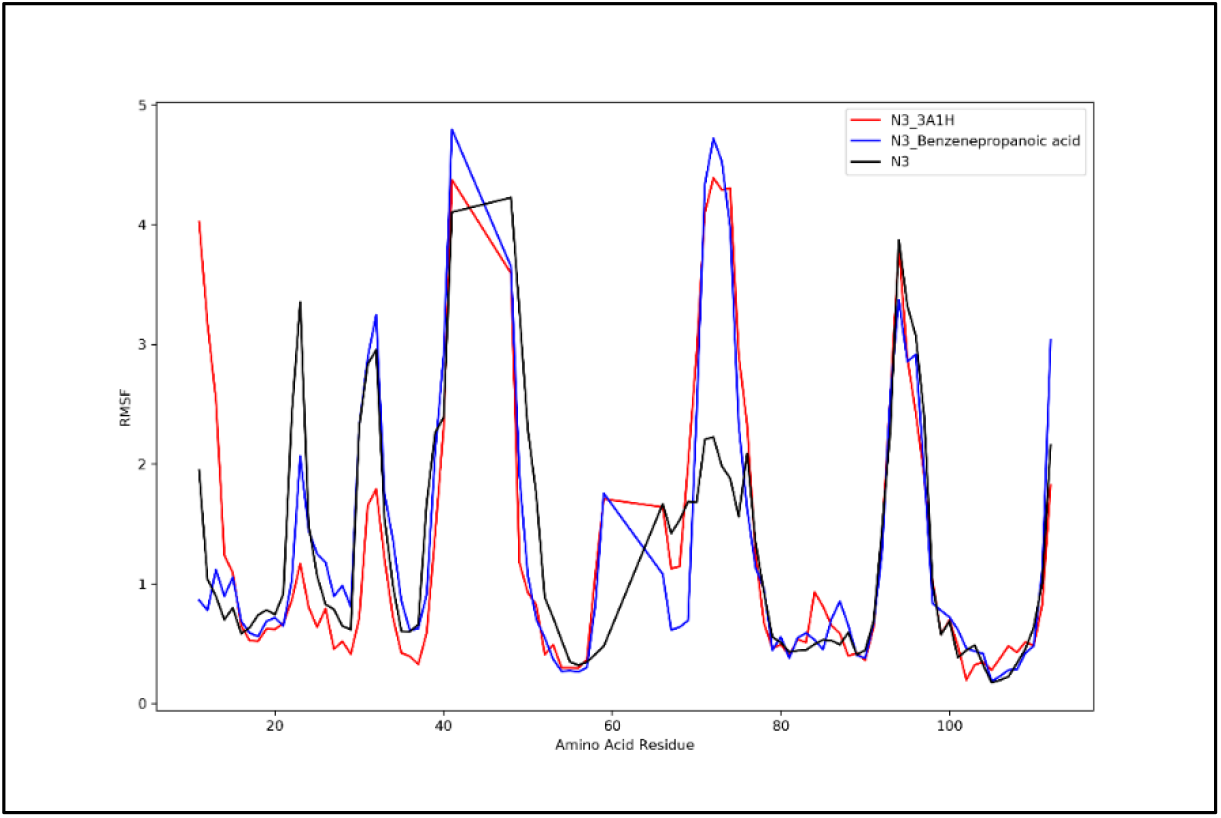
Molecular Simulation studies of N3 docked with natural ligand, Benzene propanoic acid, 3,5-bis(1,1-dimethylethyl)-4hydroxy-, methyl ester and 3A(1H)-Azulenol,2,3,4,5,8,8Ahexahydro6,8Adimethyl-3-(1-M

### Neurotrophin 4 (N4)

Molecular simulation studies of N4 with the respective phytochemicals suggest that Stigmasterol (Figure 7q) and Phytol (Figure 7o) shows better stability as compared to the binding of N4 with p75^NTR^ (Figure 7p). Looking closely at the active amino acid residue Arg98, the RMSF curves for both Stigmasterol and Phytol are lower than N4. Also, comparing the corresponding results with the PatchDock score reveals that Stigmasterol and Phytol can also potentially provide B. monnieri with properties to improve cognition and learning.

**Figure 7.**
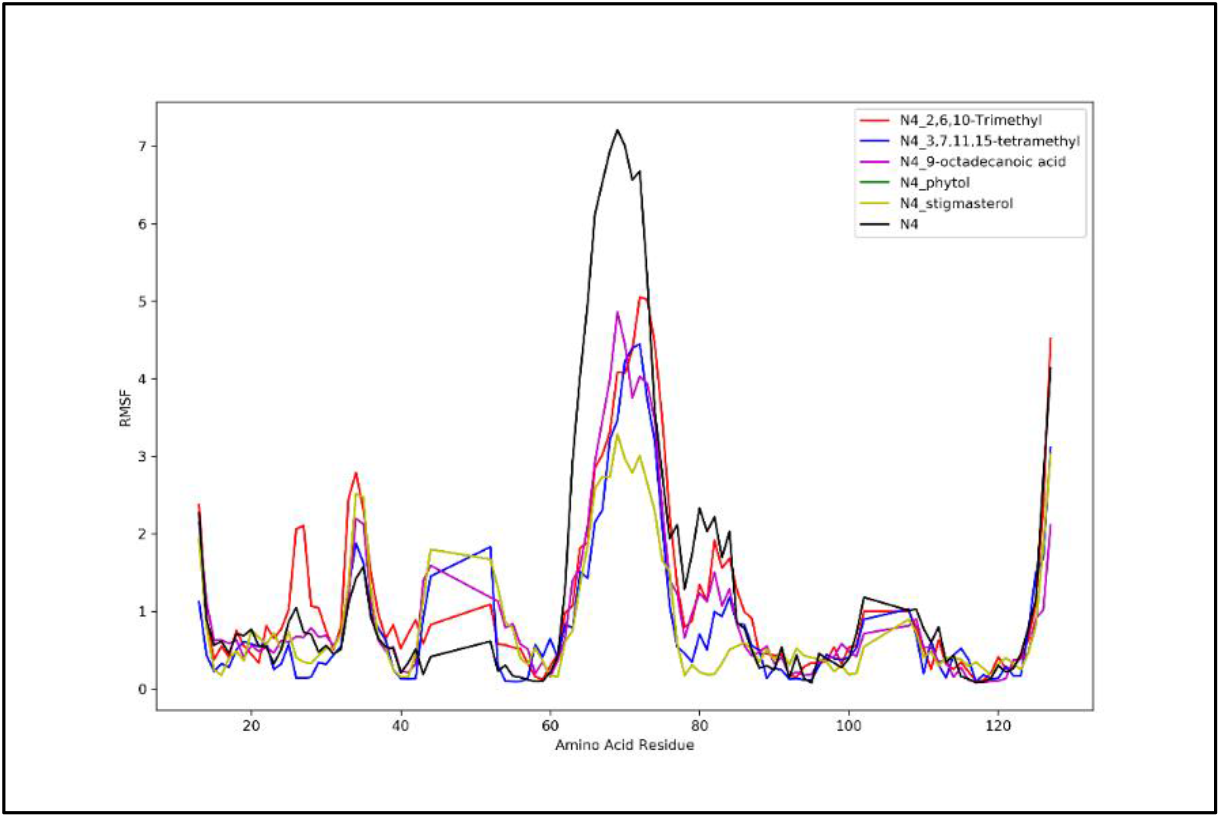
Molecular Simulation studies of N4 docked with 2,6,10-Trimethyl,14-ethylene-14-Pentadecne, 3,7,11,15tetramethyl-2Hexadecen-1-ol, 9Octadecenoic acid (Z)-, Phytol, Stigmasterol and Natural Ligand

### Nerve Growth Factor (NGF)

The molecular simulation studies of NGF with the respective phytochemical shows that Heneicosane (Figure 8c) and Nonacosane (Figure 8f) showed better results as compared with other phytochemicals. Looking at the RMSF curve of Nonacosane (dashed yellow), it can be observed that the curve at amino acid residues Trp21, Ile31 and Lys32 is below the curve of NGF (Figure 8l, black). This result suggests that the Nonacosane is more stable as compared to NGF. Also, the RMSF curve of Heneicosane (blue) at amino acid residue Lys95 is lower than the NGF curve suggesting a stable interaction. These results along with their PatchDock score suggests that Heneicosane and Nonacosane can also act as a potential source of providing B. monnieri the required anti-neurodegenerative activity.

**Figure 8.**
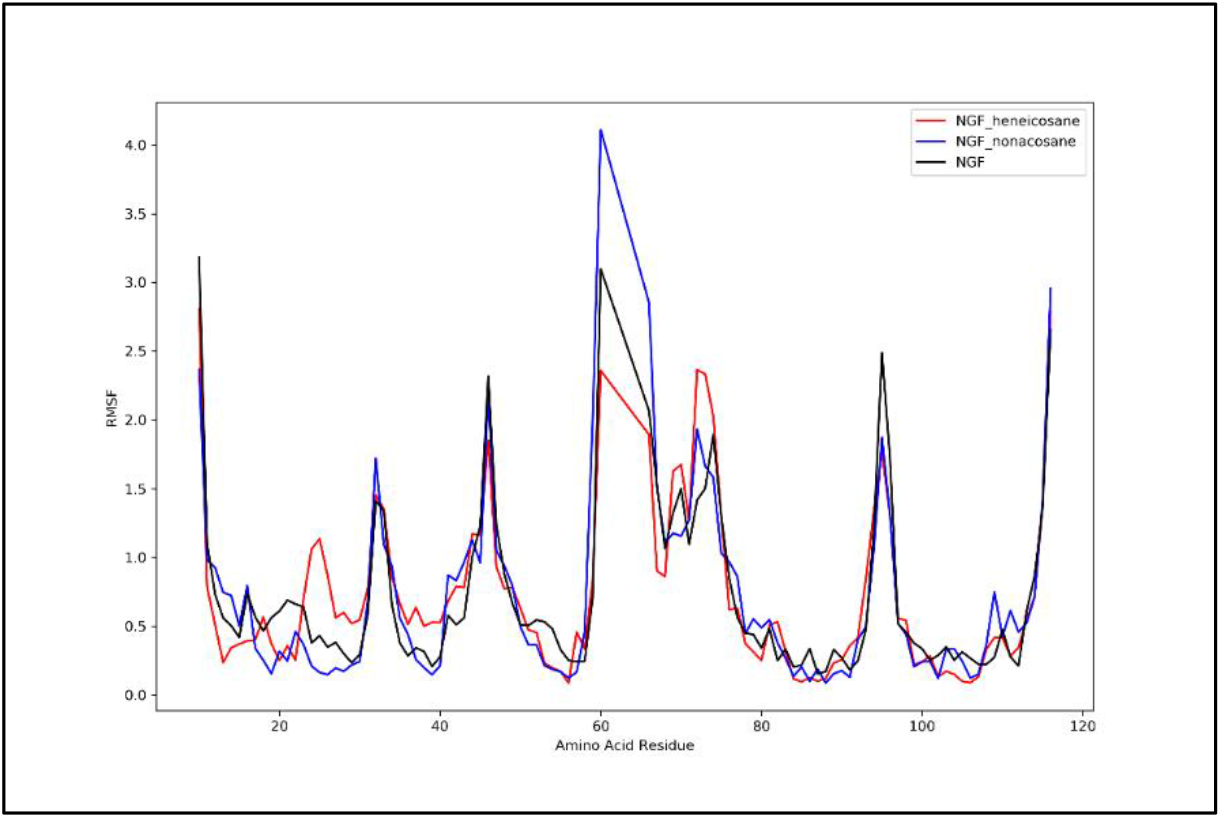
Molecular Simulation studies of NGF docked with Heneicosane, Nonacosane and Natural Ligand

### ADME Studies

Drug likeness is the property of any compound to have the potential to act as a drug. Using the SWISSADME server, we analyzed the drug likeness ability of 3 commercial drugs and the 26 phytochemicals from *B. monnieri* (Table 1). Among the drugs commercially available for the treatment of MS, Cyclophosphamide shows the maximum drug likeness by following all the drug likeness rules (Lipinski’s rule of five (RO5), Ghose filter rule, Veber rule, Egan rule and Muegge rule). The phytochemicals following all the rules of drug likeness were selected, as they correspond with Cyclophosphamide (Table 3).

The lipophilicity (LogP) is an important factor to be considered while screening compounds based on their drug likeness ability. LogP values affect the rate of absorption of drug molecules in the body. The higher the value of LogP the lower absorption of the drug molecule. The three commercial drugs have qualified almost all the drug likeness parameters. In the phytochemicals of *B. monnieri*, 1,2-Benzenedicarboxylic acid, mono(2-ethylhexyl) ester (1,2-BDA), 2-Cyclohexen-1-one, 3-(3-hydroxybutyl)-2,4,4trimethyl (Cyclo-33) and Benzenepropanoic acid, 3,5-bis(1,1-dimethylethyl)-4hydroxy-, methyl ester (BPA) showed LogP values which were higher than Cyclophosphamide suggesting that their absorption is lower as compared to the drug. But, the LogP value of 2-Cyclohexen-1-one, 4-hydroxy-3,5,5-trimethyl-4-(3oxo-1-butenyl)- (Cyclo-355) was in close proximity to that of the drug suggesting that the phytochemical has the potential to act as a natural drug molecule against MS.

The Synthetic Accessibility (SA) value of a compound suggests the ease with which the compound can be synthesized. SA value of 1 suggests that the compound can be easily synthesized whereas the value of 10 suggests that the synthesis can be challenging. The SA values of the phytochemicals were in the range of 2-3.5.

## DISCUSSION

Neurodevelopmental disorders are an important aspect to be considered. The disorders affect various parts of brain resulting in changes associated with memory, locomotion, behavior, emotions. Neurocognition and attention is an important aspect of humans to survive on a day-to-day basis. Thus, it it is of utmost importance to devise drugs that can target the neurocognitive and neurodevelopmental disorders in order to assist the patient with a better lifestyle. Synthetic drugs are not able to cure the disorders with a hundred percent certainty. Also, the risk of side effect is always associated with commercial drugs that are synthesized chemically. *Bacopa monnieri* has garnered much attention in recent years for its ability to enhance the neurocognitive and attention properties of human brain^75^. This study focuses on finding the major phytochemical that imparts *B. monnieri* with neurocognitive properties. Molecular docking studies suggest that, the medicinal property of B. monnieri is due to the combined effect of the phytochemicals of the plant rather than a single phytochemical. Further investigations with molecular dynamic simulations show that there is a stable binding of the phytochemicals of B. monnieri with Neurotrophins. The RMSF fluctuations of the phytochemicals were better than the commercial drugs available, which makes it a better drug candidate. Based on these study, four major phytochemicals were screened for ADME studies to find their characteristic likeness with that of the commercial drugs available. All the four phytochemicals followed the Drug likeness rules (Lipinski’s, Ghose, Veber, Egan, Meugge) and the bioavailability score of these phytochemicals were same as that of the commercial drug which further suggests their potential to act as a drug candidate. Future work with animal models can provide us with experimental results that will further support the current study and take us a step closer for the development of neurocognitive and neuroprotective drugs.

## CONCLUSION

*Bacopa monnieri* has been extensively used to treat the neurological disorders. Its ayurvedic roots make it a plant of medicinal purpose. It is associated with improving the brain function including intelligence and cognition^61^. *B. monnieri* extract when used to treat the animal model of Alzheimer’s (C57/Bl6 mice) showed a neuroprotective mechanism and it restricts the β-amyloid formation^74^. These results suggest that *B. monnieri* can act as a potential drug candidate against several other neurodegenerative disorders. Yet, it is unclear as to how *B. monnieri* actually causes these activities. The current work was done with aim of understanding the major involvement of phytochemical that proves *B. monnieri* to be useful in cases of neurodegenerative disorders. This study suggests that, there is no single phytochemical responsible for the *B. monnieri* activity. It is the amalgamation of majority of the natural phytochemicals of *B. monnieri* that makes it a medicinal plant that can be employed for various neurological disorders. Still, experiments will be required to find the molecular mechanism behind its action and to find out how the plant imparts its effect. Our work is a step towards unraveling the mystery behind the medicinal property of *B. monnieri*. The computational studies conducted in this work reveals the innate medicinal nature of the phytochemicals and their potential to act as a drug candidate.

